# Layered entrenchment maintains essentiality in protein-protein interactions

**DOI:** 10.1101/2024.01.18.576253

**Authors:** Luca Schulz, Jan Zarzycki, Wieland Steinchen, Georg K. A. Hochberg, Tobias J. Erb

## Abstract

Protein complexes composed of strictly essential subunits are abundant in nature and arise through the gradual complexification of ancestral precursor proteins followed by their co-evolution with the newly recruited components. Essentiality arises during co-evolution by the accumulation of changes that are tolerated in the complex state but would be deleterious for the standalone complex components. While this theoretical framework to explain how essentiality arises has been proposed long ago, it is unclear which factors cause essentiality to persist over evolutionary timescales. In this work we show that the central enzyme of photosynthesis, ribulose-1,5-bisphosphate carboxylase/oxygenase (Rubisco), rapidly started to depend on a newly recruited interaction partner through multiple, genetically distinct mechanisms that affect stability, solubility, and catalysis. We further demonstrate that layering multiple mechanisms of essentiality can lead to the persistence of essentiality, even if any given mechanism reverts through chance or selection. More broadly, our work highlights that new interaction partners can drastically re-shape which substitutions are tolerated in the proteins they are recruited into. This can lead to the rapid evolution of multi-layered essentiality through the exploration of areas of sequence space that are only accessible in the complex state.

## Main Text

Biological systems are replete with essential components. Components are essential if some process or function cannot be carried out at all if the component is missing. How any cellular component becomes essential for a living cell is an important question in evolution^1^. This is well-exemplified by protein complexes comprising several different subunits, which cannot function if any of their components are missing^2^. Such complexes develop when a novel subunit is recruited into a simpler ancestral complex, which subsequently evolves to completely depend on the new component^3–6^. This process provides an explanation of how seemingly irreducible biological complexity could have evolved. One mechanism that has been known for over two decades driving the evolution of protein complexes is that a preexisting protein starts to depend on a novel subunit if it accumulates conditionally tolerated substitutions^3,4^. Such substitutions are functionally deleterious in the absence of the novel component, but tolerated when it is present (Fig. 1A)^7,8^. Once enough such substitutions accumulated, they can make a protein-protein interaction essential because the simpler state (i.e., in the absence of the novel component) is no longer functional on its own. Empirical work has since shown that just one conditionally tolerated substitution can be sufficient to render a protein dependent on a novel component.

**Figure 1.**
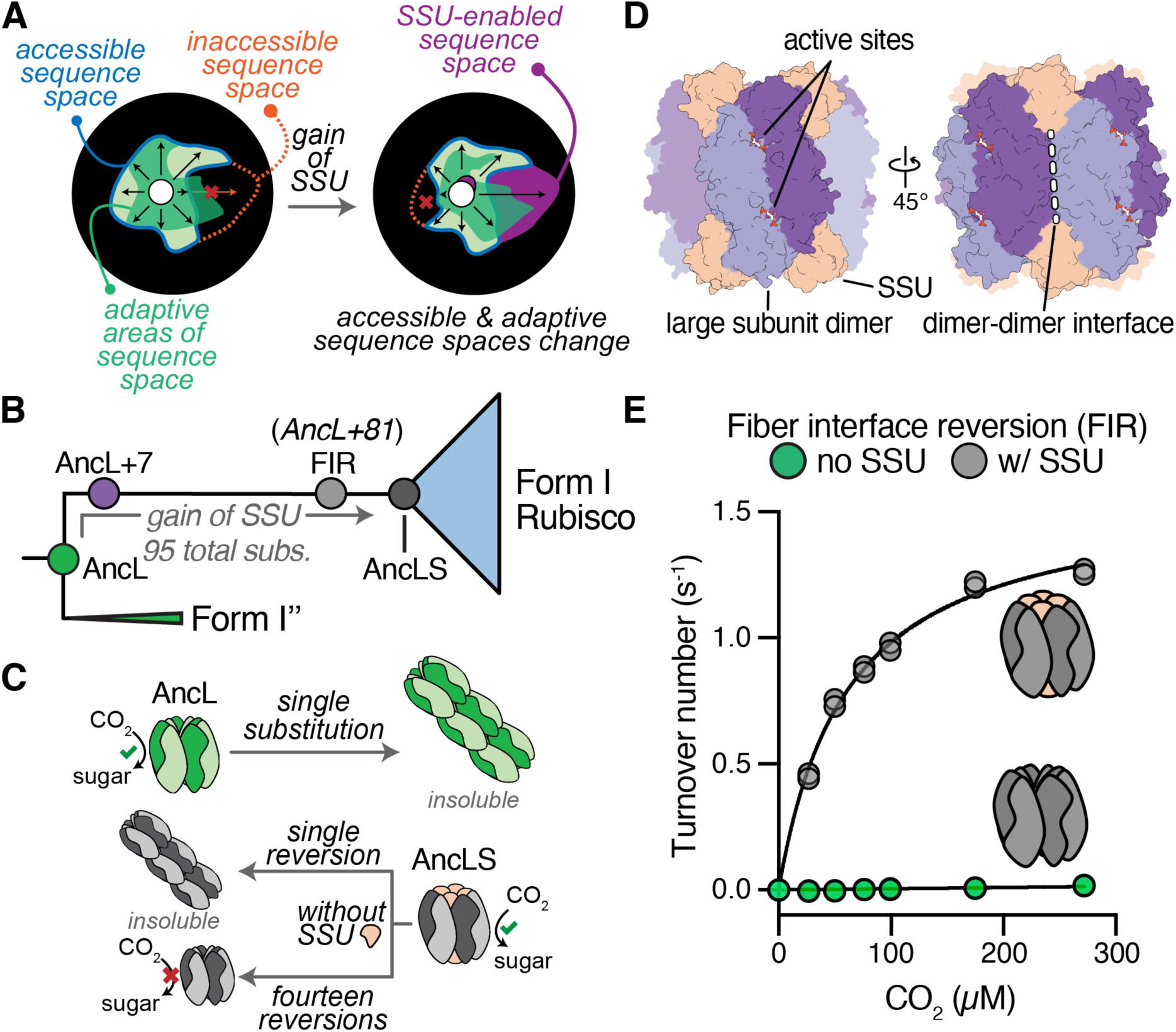
**(A)** Schematic representation of a protein’s (functional) sequence space and how the gain of complexity (here small subunit (SSU)) can modulate it. Accessible sequence space is shown in light green with blue outlines, inaccessible sequence space as an orange dotted line, functional/adaptive sequence space in green, and SSU-enabled sequence space in purple. (**B**) Schematic representation of the investigated evolutionary interval with AncL+7 and the fiber interface reversion (FIR) construct highlighted as hypothetical intermediates during the evolution from AncL to AncLS. (**C**) The ancestral AncL can become insoluble in a single historical substitution, whereas more than one reversion is required in the background of the derived AncLS to revert insolubility. Insolubility is circumvented by a combined fourteen substitution, which results in the FIR construct, which is catalytically inactive without the SSU. (**D**) AncL+7 structure surface representation highlighting the approximate position of active sites, SSU-binding sites, the dimer-dimer interface, and catalytic large subunit dimers within the overall complex. (**E**) Activity of FIR without or with addition of a ten-fold AncSSU excess under varying dissolved CO2 concentrations.

A recent example from our own work is the evolution of the CO_2_-fixing enzyme Rubisco. Modern-day Form I Rubiscos assemble into a complex of eight catalytic large subunits (LSUs) and eight non-catalytic, accessory small subunits (SSUs, L8S8 complex). Using ancestral reconstructions, we could show how Rubisco recruited the SSU for its beneficial effects on catalysis and subsequently became dependent on it through co-evolution^9^. We could recapitulate the evolution of this dependence by introducing a single historical substitution (s437W) into an ancestral Rubisco. This substitution causes Rubisco to polymerize into insoluble fibrils in the absence of the SSU. It is made harmless by binding of the SSU, which sterically prevents fibril formation, indicating that solubility (partially) drove essentiality.

The ease with which components can become essential is separate from why they remain essential. So far, why essentiality persists is not well understood^3,9–11^. If a single substitution can make a component essential, a single reversion may in principle also revert its essentiality. This is at odds with the observation that once a component becomes essential, it tends to stay essential, as is the case for Rubisco’s SSU^12–15^. One possibility is that natural selection directly acts to preserve those substitutions that made the new component essential. This would be the case if these substitutions are also directly beneficial for function of the complex. In the case of Rubisco this explanation seems unlikely: While the initial recruitment of the SSU was probably catalytically beneficial through an allosteric effect on Rubisco’s active site, substitutions that subsequently made the interaction essential, such that Rubisco could no longer function without it, apparently did not improve the enzyme at all.

Entrenchment by such functionally neutral mutations can persist if entrenching mutations are particularly likely to occur and unlikely to revert because of biases in the mutational process. We have suggested this to be the case for very hydrophobic interactions, which can become entrenched through such a mutational ratchet mechanism^3^. In many cases, however, it is not obvious that strong mutational biases exist for particular mechanisms of entrenchment. In Rubisco’s case, fiber assembly seems like a structurally delicate property that should in principle be easy to abolish by a single repulsive charge^16^. Why then do almost all extant Rubiscos continue to depend on the SSU despite billions of years of divergence?

In prior work we found that fiber assembly in a Rubisco that strictly requires the SSU for solubility (‘AncLS’) can be abolished by reverting 14 amino acid sites to their ancestral states from a Rubisco that had not yet been in contact with the SSU (‘AncL’, Fig. 1B, C & D). The resulting Rubisco variant is soluble, but catalytically inactive without the SSU (Fig. 1E). This implied the existence of a separate mechanism of essentiality that is genetically and biochemically distinct from fiber formation. Such layered entrenchment would preserve the essentiality of the SSU, even if the protein can drift in and out of any one mechanism of entrenchment.

Here we investigate the mechanism behind these deeper layers of Rubisco’s dependence on the SSU. We discover that at least two additional biochemically and genetically distinct mechanisms – stability and catalysis – can also create a dependence through a small number of historical substitutions. We find that while catalytic entrenchment is easy to create, it is very difficult to revert. Our results imply that the recruitment of the SSU dramatically changed Rubisco’s accessible sequence space, which allowed Rubisco to accumulate substitutions that are tolerated in presence of the SSU but would otherwise be deleterious. This resulted in rapid, multilayered, and genetically complex entrenchment, which has ensured the SSU’s essentiality in all Form I Rubiscos, across more than 2 billion years of evolution in diverse lineages. More broadly, our study implies that novel components can cause a drastic rewiring of a protein complex’ accessible sequence space.

## Results

### Stability entrenchment

Earlier work had shown that SSU recruitment improved Rubisco’s catalytic efficiency and enabled the accumulation of substitutions that made Rubisco depend on the SSU for catalysis^9^. Thus, we sought to find substitutions that could have caused a catalytic dependence of Rubisco on the SSU. To do this, we searched for substitutions that would induce catalytic dependence in AncL+7 (a version of AncL that carries 7 substitutions to enable its interaction with the SSU without making it essential, Fig 1C). We noticed that helix α4 (aa 261-274), which faces Rubisco’s central pore, was heavily substituted from AncL to AncLS and that three sites within that region changed to amino acids that are well conserved in Form I Rubiscos (**R**269W, **E**271R, and **L**273N). Introducing these substitutions into AncL+7 yielded a Rubisco variant (AncL+7 REL) that exhibited ∼90% decreased activity without the SSU in our assays, while activity was almost entirely recovered in the presence of AncSSU (Fig. 2A, Extended Data Figure 1). Catalytic dependence of Rubisco on the SSU can therefore be produced in few substitutions, even though the exact mechanism remained unclear at this stage.

**Figure 2.**
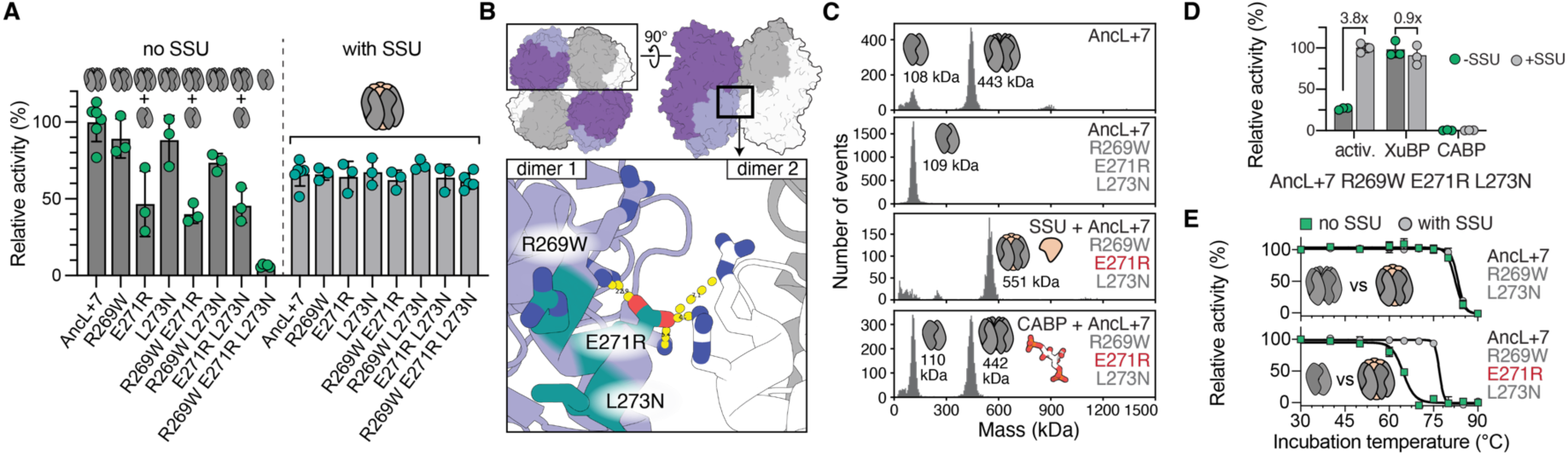
(**A**) Relative activities of AncL+7 and single, double, and triple substitution constructs on the trajectory to AncL+7 R269W E271R L273N with and without the presence of a five-fold AncSSU excess. Oligomerization state of the investigated variant at assay conditions is indicated schematically by cartoons above the bars. Activity is given relative to the activity of AncL+7 without AncSSU. (**B**) Localization of introduced substitutions in the structure of AncL+7 (PDB ID: 7QSX). Top shows a surface representation to localize the boxed cartoon representation. (**C**) Mass photometry (MP) measurements of AncL+7 and AncL+7 R269W E271R L273N (with or without AncSSU). Inferred oligomeric states are depicted as cartoons. Proteins were crash diluted from 20 µM stock concentration and measured at 50 nM final concentration, as well as MP measurement of AncL+7 R269W E271R L273N in presence of 0.2 mM CABP shows inhibitor-dependent oligomerization. (**D**) Relative activity of AncL+7 R269W E271R L273N activity assays in the presence of 1 mM XuBP or 0.2 mM CABP. Activity is given relative to the activity of the AncSSU-bound triple substitution construct in absence of inhibitory sugars. Green datapoints/dark grey bars indicate measurements in absence of AncSSU, grey datapoints/light grey bars indicate measurements in presence of a five-fold AncSSU excess. (**E**) Resistance to thermal denaturation of AncL+7 R269W L273N (remains octameric) and AncL+7 R269W E271R L273N (becomes dimeric without AncSSU), as assessed by incubating protein at varying temperatures for 1 hours, prior to assessing remaining activities at 25 °C.

We next sought to identify the biochemical basis of this dependence on the SSU. The three substitutions are localized in proximity to the Rubisco’s dimer-dimer interface. We thus hypothesized that their introduction hinders octamer formation (Fig. 2B). Indeed, instead of forming an octamer, the triple mutant was dimeric in the absence of AncSSU, as determined by mass photometry (MP, Fig. 2C). Even a single substitution (E271R) was sufficient to abolish octamer formation (Extended Data Figure 2) and led to ∼60% reduced activity under assay conditions, whereas activity in presence of AncSSU was unchanged (Fig. 2A). The addition of AncSSU recovered octamerization and led to the formation of canonical L8S8 complexes (Fig. 2C). This indicates that the SSU compensates for the detrimental effect of these substitutions by holding together these weaker octamers. Overall, this data suggested that the SSU buffers substitution at the dimer-dimer interface, which destabilize oligomerization.

However, it is unclear if there is a functional benefit associated with octamer formation in Form I Rubisco and recent studies suggest that oligomerization is not crucial for activity in Form II Rubiscos^17^, which acquired their higher order assemblies (dimers, tetramers, and hexamers) independently from Form I Rubiscos. We thus tested if AncSSU-dependence arose because octamers were crucial for function in Form I Rubiscos. While we could not observe substrate-dependent (RuBP) oligomerization, which has been reported for some Rubiscos^18^, we noticed that the addition of known active site inhibitors of Rubisco, namely xylulose-1,5-bisphosphate (XuBP)^19^ or 2-carboxyarabinitol-bisphosphate (CABP)^20^, induced octamer formation in AncL+7 REL (Fig. 2C and Extended Data Figure 3). We additionally solved the crystal structures of AncL+7 REL in CABP-bound forms (with and without AncSSU), in which the protein was present exclusively as an octamer with or without AncSSU (PDB 8QMW and 8QMV, respectively). Thus, a strong enough active site binder or very high concentrations of the protein can force AncL+7 REL into octamers, even though its native substrate RuBP cannot (Extended Data Figure 4). This implies that there is a direct link between Rubiscos active site and the dimer-dimer interface and conversely suggests, that Form I Rubiscos need to be octamers to be fully catalytically active.

**Figure 3:**
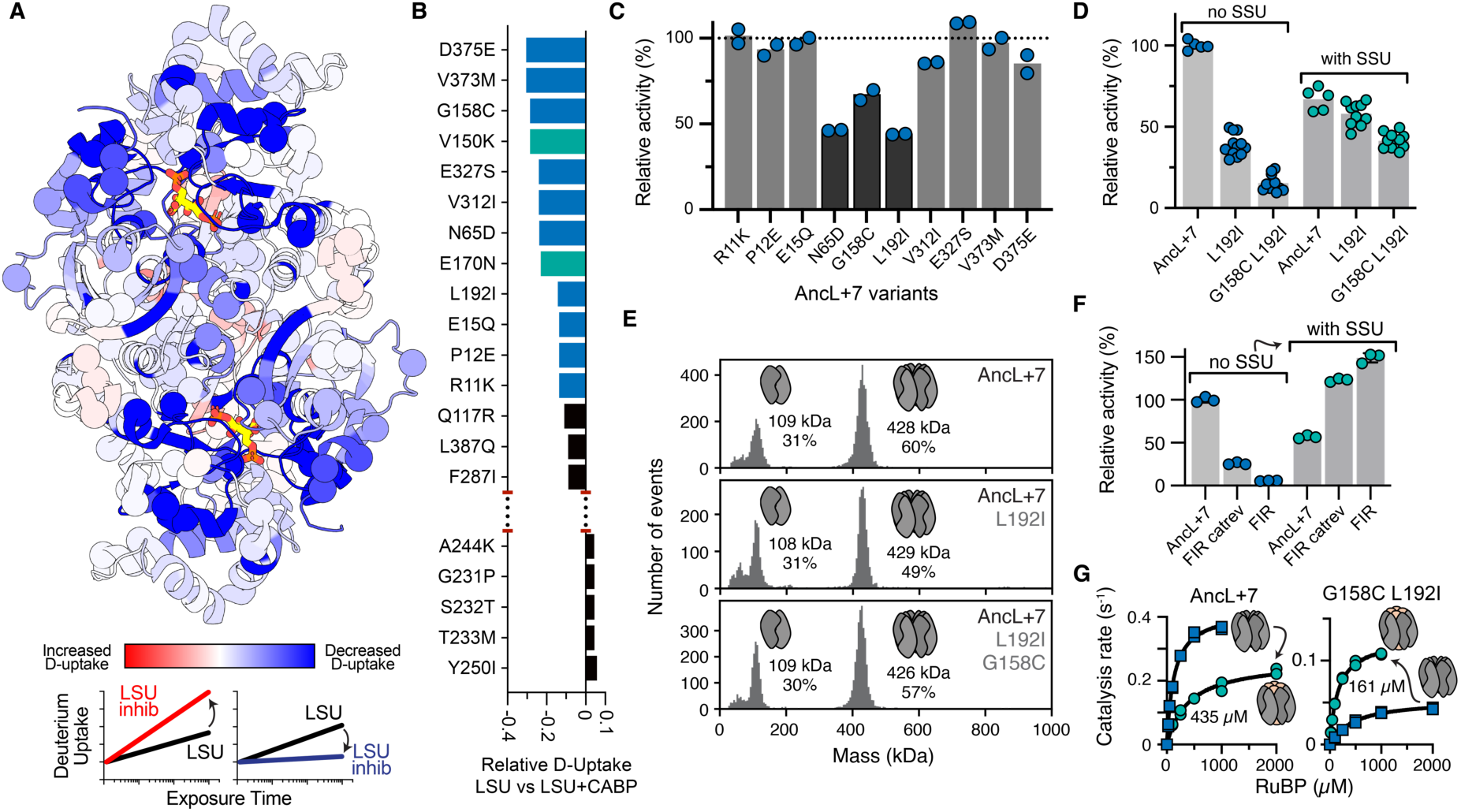
(**A**) Relative hydrogen-deuterium exchange mass spectrometry (HDX) deuterium uptake rates of AncL+7 with versus without inhibitory sugar (CABP) measured 95 s after protein dilution into deuterated buffer and projected onto a catalytic dimer of AncL+7 (PDB ID: 7QSX). Scheme at bottom indicates the expected deuterium uptake behavior of regions with increased and decreased uptake rates. (**B**) Substituted sites with the highest and lowest relative D-uptake rate differences when comparing AncL+7 versus AncL+7 with CABP in ranked form. Investigated substitutions from AncL to AncLS are indicated in blue (this study) or teal (previous work^9^). (**C**) Relative activity of AncL+7-based single substitution constructs, relative to the activity of AncL+7 without AncSSU. (**D**) Relative activity of AncL+7 single and double substitution constructs with and without presence of a five-fold AncSSU excess. Activity is relative to the activity of AncL+7 without AncSSU. Measured oligomerization states are annotated in cartoon form above the respective bars. (**E**) Mass photometry measurements of AncL+7 and relevant single or double substitution constructs. Inferred oligomeric state is shown schematically, measured mass is listed, and the total percentage of binned events relative to all events of the measurement are listed. (**F**) Relative activity of AncL+7 and the fiber interface reversion (FIR) construct, as well as the FIR construct with “G158C” and “L192I” reverted to their state in AncL+7 (catrev) measured in absence (blue) and presence (teal) of a tenfold AncSSU excess. (**G**) RuBP Michaelis Menten kinetics for AncL+7 and AncL+7 G158C L192I without (blue) and with (teal) presence of a five-fold AncSSU excess. Inset values list the measured *Km*(RuBP) in presence of AncSSU.

We were able to test this theory directly because unlike with plant Rubiscos, XuBP does not inhibit catalysis in AncL+7-derived Rubiscos (Extended Data Figure 5). This enabled us to measure activity in presence of XuBP while artificially shifting Rubisco’s oligomeric state towards the octamer. Indeed, adding 1 mM XuBP to activity assays of AncL+7 REL containing 3 mM RuBP recovered activity in the absence of AncSSU (Fig. 2D). This suggested that XuBP can still efficiently diffuse out of Rubisco’s active site after inducing oligomerization, which then allows for binding and turnover of RuBP and CO_2_ before the octamer dissociates. Notably, the effect was XuBP-specific and could not be induced by non- or mono-phosphorylated xylulose or the tight-binding active site inhibitor CABP, which stays bound and inhibits catalysis (Extended Data Figure 5B & 6A). To independently test for octamer-dependence of catalysis, we crash diluted Rubisco from high protein concentrations, at which we hypothesized octamers to still be formed (>100 µM), to a normalized assay concentration. This resulted in recovered activity when initiating assays from high pre-dilution concentrations (Extended Data Figure 6B). Together these results imply that Form I Rubiscos are only active as octamers. This allows for entrenchment of the SSU, which links together two adjacent LSU dimers. This bridging provides extra stability to the interaction between LSU dimers and allows the dimer-dimer interface itself to be subsequently degraded again through substitutions in this region.

Lastly, we noticed that AncL+7 REL, as well as all other constructs that form dimers instead of octamers without AncSSU, were drastically destabilized towards thermal denaturation (∼30 °C), which was partially recovered by AncSSU addition (Fig. 2E, Extended Data Figure 7), implying that LSUs in dimers are less stable than LSUs associated into octamers. In hot environments, this could potentially serve as separate mechanism of entrenchment independent of octamer-dependent catalysis. Increased stability as an octamer could plausibly be a direct selective advantage given the right environment, or could offer evolution room for adaptive, yet destabilizing evolution^21,22^.

### Catalytic entrenchment

Our observations on AncL+7 REL imply that the SSU could become entrenched because it provided stability to octamers. However, this cannot be the sole reason for AncLS’ catalytic dependence on AncSSU: our fiber interface reversion construct of AncLS (which led to this investigation) was octameric, yet catalytically inactive without SSU (Fig. 1E). This implied that there must be yet another mechanism for catalytic entrenchment, genetically and structurally independent from the ability to assemble octamers.

We could not rationally identify an obvious set of candidate substitutions for catalytic SSU-dependence from the sequence differences between AncL and AncLS. We therefore sought an alternative way to identify plausible sites. Based on our findings regarding higher oligomerization states in the presence of XuBP and CABP we reasoned that sites distal from the active site that undergo changes upon substrate binding would be promising candidates in which substitutions could lead to SSU dependence ^23^.

To find such sites, we performed hydrogen/deuterium exchange mass spectrometry (HDX) on AncL+7 alone and bound to the transition state analogue CABP (Supplementary Data, Extended Data Figure 8). HDX quantifies the time-dependent exchange of hydrogen to deuterium in proteins, which is dependent on solvent accessibility. When comparing conditions, differences in solvent accessibility can stem from structural re-arrangements or changes in protein motion. As expected, comparing HDX rates between the two conditions revealed ordering of the active site and regions known to be involved in catalysis (e.g., loop 6^24^ and the 60s loop^25^) upon inhibitor binding. It also revealed regions up to ∼15 Å from the active site that decrease their deuterium uptake upon inhibitor binding (Fig. 3A).

We chose to investigate the effect of ten substitutions in regions with the most drastic HDX response to inhibitor binding (Fig. 3B). When introduced into AncL+7 as single substitution, three of the ten substitutions led to a 40-60% reduction in activity, relative to AncL+7 (Fig. 3C). This included substitutions such as an internal leucine to isoleucine (L192I), which reduced activity by ∼60%. Activity of this construct could largely be rescued by addition of AncSSU (Fig. 3D). When further combined with G158C, a second substitution that had a deleterious effect on catalysis in absence of the SSU, activity was reduced to ∼25% yet could still largely be rescued by AncSSU addition (Fig. 3D). Both the AncL+7 L192I and the AncL+7 G158C L192I mutants retained their octameric assembly, indicating that a mechanism distinct to the loss of oligomerization was causing dependence on the SSU for catalysis (Fig. 3E). Reverting C158G and I192L in the soluble, yet catalytically inactive, fiber interface reversion construct recovered some catalytic competence (Fig. 3F). This indicated that C158G and I192L were only partially responsible for Rubisco’s catalytic entrenchment and that additional substitutions deepened Rubisco’s catalytic dependence. Like the other forms of entrenchment, catalytic entrenchment can thus be created in a small number of substitutions but requires a larger set of reversions to abolish.

We next investigated whether G158C L192I substitutions were functionally neutral, in the same way that the fiber entrenching substitution was. In AncL+7 the *K*_m_(RuBP) increased roughly 3-fold upon AncSSU addition, whereas this effect was inverted in AncL+7 G158C L192I and *K*_m_(RuBP) decreased from 435 µM to 161 µM (Fig. 3G). While this positive effect still coincided with an overall decreased catalytic rate, it may be beneficial, given the right conditions. This is circumstantial evidence that conditionally tolerated substitutions can be advantageous, implying that novel subunits indeed open adaptive evolutionary paths that would otherwise not be accessible.

## Discussion

Using Rubisco’s dependence on the SSU as an example, we demonstrate a potentially very general principle that explains the persistence of essentiality, even in the absence of strong mutational biases favoring particular mechanisms of entrenchment: Novel interactions can apparently drastically change the accessible sequence space of proteins, opening up many new genotypes that are only accessible in the presence of the new component. In the case of Rubisco, this effect also encompassed positions that were far from the SSU binding site, which then led to rapid and multilayered entrenchment of the new interaction. We can currently only speculate about the biochemical mechanism behind this effect. Crystal structures of AncL+7 with and without the SSU show only minor differences in Rubisco’s ground state structure^9,24^. We therefore believe that SSU binding alters Rubisco’s dynamic motion on functionally relevant timescales across the protein, which in turn changes the effect of substitutions at sites with modified dynamics. Future work will have to test this prediction empirically.

Our work shows that each mechanism of entrenchment we discovered can be established through a small number of substitutions in AncL, but reverting the corresponding substitutions in AncLS is not enough to yield an SSU-independent, fully catalytically active Rubisco. This implies that there must be more entrenching states and likely additional mechanisms that remain to be discovered. One problem is that we do not know the temporal order of the substitutions that separate AncL from AncLS. Whether or not certain substitutions cause dependence on the SSU relies on their exact sequence context. For example, entrenchment by abolishing oligomerization seems context dependent in this way: the fiber interface reversion construct contains all states that when introduced into AncL+7 make it fall apart into a dimer, however, it nonetheless forms octamers in the absence of the SSU. Still, we have been able to alleviate entrenchment of solubility and catalysis either fully or partially, in AncLS by reverting substitutions that were causal when introduced into AncL. This gives us confidence that the genetic and structural mechanisms we have discovered were relevant for Rubisco’s historical evolution.

Form I Rubiscos have remained dependent on the SSU over more than 2 billion years of evolution, even though selection presumably does not act directly to maintain the SSU’s essentiality. The multiplicity and depth of the entrenchment we have discovered in our ancestors explains why: it ensures that the SSU stays essential even if Rubisco drifts out of any one entrenching mechanism. This has happened at least once, in a cyanobacterial Rubisco that is partially soluble without the SSU (and hence has reduced its entrenchment via fiber formation) but is barely active in the absence of the SSU^26^.

The SSU is not the only novel component Form I Rubiscos recruited in their history. Several dedicated assembly chaperones became completely essential for Rubiscos within the plant lineage^27^. Along the same lineage, Rubiscos additionally started to become dependent on Rubisco activases^28,29^ – dedicated ATPases that remove inhibitory sugar phosphates from its active site. It is tempting to speculate that each addition enabled Rubisco to access new catalytic optima that reside in previously inaccessible parts of sequence space, which were opened up by the addition of a novel component^30^. Alternatively, it is possible that Rubisco became addicted to such components without any functional benefit, solely because they unlocked a vast enough area of sequence space from which Rubisco has since simply failed to escape.

## References and Notes

## Supporting information

HDX-MS source data

## Acknowledgements

The X-ray diffraction data was collected at the P14 beamline of the PETRA III storage ring (DESY, Hamburg, Germany). We would like to thank Selina Storm for the assistance at the beamline. The authors are grateful for generous support from the Max Planck Society. L.S. thanks the Joachim Herz Foundation for support in form of an Add-On fellowship for Interdisciplinary Life Sciences.

## Author Contributions

L.S., T.J.E., and G.K.A.H. conceived the project, analyzed data, and planned experiments. L.S. performed molecular work, phylogenetics, protein purification, enzyme kinetic analysis and specificity constant measurements, x-ray analysis, comparative in vitro biochemistry, and MP measurements. W.S. performed and analyzed HDX-MS experiments. J.Z. collected, solved, refined, and analyzed x-ray structures. T.J.E., and G.K.A.H. supervised the project. L.S., T.J.E., and G.K.A.H. wrote the manuscript with contributions and comments from all authors.

## Data availability statement

Raw data will be deposited on Edmond, the Open Research Data Repository of the Max Planck Society for public access. The atomic structures reported in this paper were deposited to the Protein Data Bank in Europe under accession codes 8QMV and 8QMW.

## Methods

### Chemicals and reagents

Chemicals were of the highest commercially available purity from Sigma-Aldrich and Carl Roth. NaH^14^CO_3_, K^14^CN, and D-[2-^3^H] glucose was obtained from Hartmann Analytics (Germany). Biochemicals (for cloning and protein production/purification) were obtained from Thermo Fisher Scientific, Macherey-Nagel, and New England Biolabs. 3-phosphoglyceric phosphokinase and glyceraldehyde 3-phosphate dehydrogenase were obtained from Sigma-Aldrich.

Rubisco-specific chemicals (D-Ribulose-1,5-bisphosphate (RuBP), Xylulose-1,5-bisphosphate (XuBP), 2-carboxyarabinitol-1,5-bisphosphate (CABP), and [1-^3^H] RuBP) were synthesized following published protocols^31–33^ as described below.

Non-radioactive RuBP was synthesized from D-Ribose-5-phosphate disodium salt (Sigma #R7750, 1.28g) in a 400 mL reaction containing 40 mM MgCl2, 0.25 mM ATP, 15.2 mM creatine phosphate, 5 mM dithiothreitol, 6.5 mg creatine phosphokinase (Sigma #CK-RO), 6 mg *Arabidopsis thaliana* phosphoriboisomerase, and 4.6 mg *Synechocystis* sp. PCC 6803 phosphoribokinase. RuBP was purified using an XK 50/70 AG1-X8(Cl) column equilibrated to 3 mM HCl, 100 mM NaCl and eluted with a 6.25 column volume gradient increasing in salt concentration to 3 mM HCl, 250 mM NaCl. Collected fractions were lyophilized until dry, stored at -80 °C until further use, and subsequently dissolved in 50 mL of 3 mM HCl. The dissolved sample was desalted using a Sephadex G-10 XK26/100 column equilibrated and operated with 3 mM HCl. Collected fractions were pooled, aliquoted, and used as directly after determining the RuBP concentration using a photospectrometric coupled assays described below.

XuBP was synthesized by aldol addition of dihydroxyacetone phosphate to glycoaldehyde phosphate catalyzed by rabbit muscle aldolase (Sigma #A2714). Glycoaldehyde phosphate barium salt was synthesized in an aqueous 150 mL reaction by partially dissolving 12 mmol (2.33 g) glycerol phosphate magnesium salt hydrate (Sigma #17766) and 11 mmol sodium meta-periodate (2.35 g). The pH of the solution was adjusted to 6.0 with 1 M HCl followed by incubation at 37 °C for 1 hour. Afterwards 2 mmol glycerol was added to quench the reaction and the pH was adjusted to 7 with 1 M NaOH. 22 mmol (4.58 g) barium chloride was added and the reaction was incubated on ice for 1.5 hours. Precipitated BaIO_3_ was removed by filtration through a 0.22 µm sterile filter, the filtrate was mixed with four volumes (600 mL) absolute ethanol and stirred for 15 min to precipitate the glycoaldehyde phosphate barium salt. The precipitate was collected by centrifugation at 5000 x g for 20 min at 25 °C and subsequently dried by lyophilization.

For the aldolase-catalyzed synthesis of XuBP, 52.3 mg glycoaldehyde phosphate barium salt and 24 mg dihydroxyacetone phosphate (Sigma #37442) were partially dissolved in 2.5 mL 10 mM MES-NaOH pH 6.5 and 30 U rabbit muscle aldolase was added to initiate the reaction. The reaction was incubated at 37 °C for 3 hours. Over the course of the reaction a white precipitate of XuBP-Ba_2_ formed. After reaction completion, the XuBP-Ba_2_ precipitate was collected by centrifugation at 3200 x g for 15 minutes. The visibly distinct white XuBP-Ba_2_ precipitate was separated from precipitate that was present from the onset of the reaction.

The white XuBP-Ba_2_ precipitate, as well as the soluble reaction supernatant, were acidified to pH 3.0 via the slow addition of activated AGW50X-HCl-resin (∼120 µL), which dissolved the precipitate. The resin was removed by filtration through a 0.45 µm syringe filter and the syringe was washed with 1 mL 3 mM HCl. The soluble XuBP filtrate was further purified by separation on a HiPrep Q HP 16/10 column equilibrated to 3 mM HCl with a gradient to 3 mM HCl, 350 mM NaCl over 10 column volumes (elution at ∼125 mM NaCl). Fractions were assayed for the presence of XuBP using a photospectrometric coupled assay based on rabbit muscle aldolase and NADH oxidation via the activity of glycerol phosphate dehydrogenase. Fractions were pooled, the XuBP concentration was determined using a photospectrometric depletion assay, and fractions were stored at -80 °C until further use.

[1-^3^H] RuBP was synthesized enzymatically from D-[2-^3^H] glucose. Reactions were assembled in 1.7 mL containing 33 mM bis-tris-propane-HCl pH 7.5, 10 mM MgCl_2_, 0.5 mM ATP, 2 mM creatine phosphate, 1.5 mM NADP, 10 mM dithiothreitol, 34 µg creatine kinase (Sigma #10127566001), 6.8 U Hexokinase (Sigma #1142636001), 6.8 U glucose-6-phosphate dehydrogenase (Sigma #G5760), 6.8 U 6-phosphate gluconate dehydrogenase (Sigma #P4553), 0.85 mg phosphoribulokinase from *Synechocystis* sp. PCC6803, 250 µL ^3^H-glucose (evaporated over night at room temperature to a volume of ∼25 µL), and 0.6 mM glucose (non-radioactive). Reactions were incubated for 4 hours at 25 °C, 300 rpm shaking in a tabletop shaker. Subsequently 250 µL crude reaction aliquots were stored at -80 °C and used as such for specificity assays.

^14^C-containing CABP was synthesized by using 1 mCi K^14^CN per synthesis reaction. At 55.5 mCi/mmol, 18.2 µmol radioactive K^14^CN was combined with 14.6 µmol RuBP. To this end, 431 µL of a 36 mM RuBP stock dissolved in 3 mM HCl was diluted with 200 µL 1 M tris-acetate pH 8.5 and the pH was adjusted to 8.3 using 1 M tris (free base). The buffered RuBP was added to the K^14^CN vial and left to react at room temperature for ∼48 hours. Crude reactions were purified via gravity Dowex 50W-X8(H^+^) columns with 1.8 mL beads as the bed volume. Resins were washed and equilibrated with 10 mL distilled H_2_O prior to application of the reaction to the resin. The flow-through, as well as 2x 1 mL ddH_2_O elution fractions, were collected, pooled, and subsequently dried under a gentle stream of N_2_. Dried reaction product was dissolved in 8 mL of 50 mM Bicine-NaOH pH 9.3, aliquoted, and the radioactivity/specific activity was determined by counting aliquots/dilutions of the synthesized carboxypentinol-1,5-bisphosphate mixture. Fresh aliquots of CPBP were incubated overnight at 4 °C prior to first use to allow for de-lactonization.

### Molecular cloning and vector construction

All primers were obtained from Eurofins genomics. A list of all primers is provided in Supplementary Table 1. Constructs destined to carry an N-terminal His-tag were cloned into the standard *E. coli* expression vector pET16b (Merck Chemicals) and those destined to carry a C-terminal His-tag were clonsed into the standard *E. coli* expression vector pET28b (Merck Chemicals). Genes were amplified using 2x Phusion High-Fidelity PCR Master Mix (Thermo Fisher) with primers that introduce complementary overhangs to the respective PCR-linearized vectors. PCR products were used to construct the desired vectors using home-made 1.33x Gibson assembly master mix^34^. Assembly was verified by DNA sequencing (MicroSynth). All plasmids in this study are listed in Supplementary Table 2.

Single-site mutants, insertions, truncations, and deletions were created using an adapted Q5 Site-Directed Mutagenesis procedure. Primers were designed using NEBasechanger (nebasechanger.neb.com) and used to amplify the entire desired vector with 2x Phusion High-Fidelity PCR Master Mix. PCR products (0.5 µL straight from the PCR) were used in reactions containing 2.5 µL ddH_2_O, 0.5 µL T4 DNA Ligase (NEB #M0202L), 0.5 µL T4 Polynucleotide Kinase (NEB #M0201L), and 0.5 µL DpnI (NEB #R0176L) for 2 hours at 25 °C. Subsequently the entire reaction was transformed into NEB Turbo Competent *E. coli* (NEB #C2984H, list of strains in Supplementary Table 3) before the resulting vector was purified and mutagenesis success was verified by sequencing (MicroSynth).

### Protein production and purification

To produce proteins, plasmids encoding the respective genes were transformed into chemically competent *E. coli* BL21(DE3) cells and grown overnight on selective LB agar plates containing either 100 µg mL^-1^ ampicillin or 50 µg mL^-1^ kanamycin at 37 °C overnight. Grown colonies were used to inoculate expression cultures in terrific broth (TB) medium containing the respective antibiotics. Expression cultures were grown in a shaking incubator at 37 °C to an OD_600_ of 0.5-1.0, cooled down to 25 °C, induced with 0.5 mM isopropyl-β-D-thiogalactoside (IPTG), and subsequently left to produce protein overnight.

Cells were harvested by centrifugation at 8,000 x g for 10 minutes at 10 °C and cell pellets that were not used immediately were stored at -20 °C until further use. For purification, cell pellets were re-suspended in buffer A (50 mM HEPES-NaOH, 500 mM NaCl, pH 7.6). Resulting suspensions were lysed using a Sonopuls GM200 sonicator (BANDELIN Electronic) at an amplitude of 55% with three consecutive cycles of 30 pulses over 60 s and the crude lysates were clarified by centrifugation at 13,300 x g and 4 °C for 1 hour. Clarified lysates were filtered (0.45 µm syringe tip filters) and applied to pre-equilibrated Protino Ni-NTA Agarose (Macherey-Nagel) beats in a gravety column. After loading the resin was washed with 45 column volumes of 15% (v/v) buffer B (50 mM HEPES-NaOH, 500 mM NaCl, 500 mM imidazole, pH 7.6) in buffer A before protein was eluted with 8 column volumes of 100% buffer B. The eluate was concentrated to a total volume of 2.5 mL and desalted using PD-10 desalting columns (GE Healthcare) and desalting buffer (25 mM Tricine-NaOH, 75 mM NaCl, pH 8.0). For crystallization and large-scale purifications, protein was further purified via size exclusion chromatography (SEC) on a Superdex 200 pg, HiLoad16/600 column (GE Healthcare). Elution fractions containing pure protein were determined via SDS-PAGE analysis on a 4-20% gradient gel (Bio-Rad), pooled, and concentrated using 50 kDa MWCO centrifugal filters (Amicon). Purified Rubiscos in desalting buffer were used for crystallization immediately or stored at -20°C in desalting buffer until further use. For crystallization, a ∼2-fold molar excess of CABP (dissolved in 100 mM Bicine, 17.6 mM MgCl_2_, pH 8.0) relative to the Rubisco concentration (determined by absorption at 280 nm) was added. Rubisco was carbamylated in a 3% (v/v) CO_2_ atmosphere at 30 °C for 1 hour and used for setting crystal plates immediately (see “crystallization and structure determination”).

### Thermal stability assay

Apparent melting temperatures of Rubisco were determined as published previously^35,36^. In short, 20 µM purified Rubisco was activated in 50 mM HEPES-NaOH (pH 8.0), 20 mM MgCl_2_, 40 mM NaHCO_3_, and 0.02 mg/mL carbonic anhydrase from bovine erythrocytes (Sigma-Aldrich) for 15 min at 25 °C. Activated Rubisco was incubated for 60 min at varying temperatures from 30 – 90 °C prior to a 5 min incubation on ice and a 1:10 dilution into assay mixtures. Activity assays contained a final concentration of 100 mM HEPES-NaOH (pH 8.0), 0.8 mM NADH, 10 mM MgCl_2_, 0.5 µM 3-phosphoglyceric phosphokinase (pgk), 0.5 µM glyceraldehyde 3-phosphate dehydrogenase (gapdh), 5 mM ATP, 45 mM NaHCO_3_, 0.02 mg/mL carbonic anhydrase and 10x diluted activation mixture (∼2 µM activated Rubisco). Assays were initiated by the addition of 2 mM RuBP and reaction progress was followed in a microplate reader (Tecan) at 25 °C by measuring the consumption of NADH, as determined by decreasing absorbance at 340 nm. Activities were normalized to the activity of Rubisco incubated at 30 °C.

### Radiometric kinetic analysis and CO_2_/O2 specificty assays

^14^CO_2_ fixation assays were carried out at 25 °C in 0.5 mL filled into 7.7 mL septum-capped glass scintillation vials^37^. Assay buffer (100 mM EPPS-NaOH (pH 8.00), 20 mM MgCl_2_, 1 mM) and other required components were pre-equilibrated with CO_2_-free N_2_ gas. Beyond assay buffer, reactions contained an additional 0.01 mg/mL carbonic anhydrase, 2.2 mM self-synthesized RuBP and 5-70 mM NaH^14^CO_3_ (corresponding to 100 to 800 µM ^14^CO_2_). Dissolved CO_2_ concentrations were calculated using the Henderson-Hasselbalch equation with pK values for carbonic acid of pKa1 = 6.25 and pKa2 = 10.33, while accounting for assay volume and headspace volume. Active site contents of purified Rubiscos were quantified for each data set by performing [^14^C]-2-CABP binding assays on 0.2 nmol purified Rubisco, followed by separation from free ligand by size exclusion chromatography^38^ (Isera SEAgel-Column, custom made for HPLC) and active site quantification by scintillation counting. Purified Rubisco (∼5-30 µM active sites) was activated in assay buffer supplemented with 50 mM NaHCO_3_ and 20 µL of the activation mixture was used to initiate the assay. Reactions were carried out at 25 °C for 2 min before quenching with 200 µL 50% (v/v) formic acid. The specific activity of NaH^14^CO_3_ was determined by completely turning over 22.6 nmol RuBP using the highest employed NaH^14^CO_3_ concentration in a 60 min reaction.

Specificity assays were performed as published previously with slight adaptations^32,37^. In short, purified Rubisco was incubated in 20 mL septum capped glass scintillation vials containing 1 mL 30 mM triethanolamine (pH 8.30), 15 mM Mg-acetate, and 0.01 mg/mL carbonic anhydrase. Assays were equilibrated in defined gas mixtures containing either 995,000 ppm O_2_ and 5000 ppm CO_2_ (for low specificity variants) or 999,291 ppm O_2_ and 709 ppm CO_2_ (for high specificity variants, Air Liquide, Germany) before starting reactions by addition of [1-^3^H]-RuBP. Reactions were incubated for 60 minutes under constant gassing and subsequently dephosphorylated using alkaline phosphatase (10 U/reaction), separated on an HPX-87H column (Bio-Rad), and relative amounts of glycerate and glycolate quantified by flow-through scintillation counting. Specificity was calculated as described previously^32^.

### Ribulose-1,5-bisphosphate Michaelis-Menten kinetics

To determine the *K*_M_(RuBP), photospectrometric reactions were set up with slight adaptations to previously published protocols^38^. Reactions were carried out in 100 mM HEPES-KOH (pH 8.00) and contained 10 mM MgCl_2_, 2.5 mM ATP, 0.3 mM NADH, 2.5 U/mL pgk, 5 U/mL gapdh, 0.02 mg/mL carbonic anhydrase, 55 mM NaHCO_3_, varying amounts of RuBP (0 – 5 mM), 0.2-2 µM activated Rubisco, and a 5-fold SSU excess, if applicable. Rubisco was activated by a 30-minute pre-incubation in 50 mM HEPES-KOH (pH 8.00), 10 mM MgCl_2_, 20 mM NaHCO_3_, and 0.02 mg/mL carbonic anhydrase. Reaction progress was followed by measuring the depletion of NADH at 340 nm (Abs_340_).

### Mass photometry

Mass photometry measurements were carried out on microscope coverslips (1.5 H, 24 x 50 mm, Carl Roth) with CultureWell^TM^ Reusable Gaskets (CW-50R-1.0,50-3mm diameter x 1 mm depth) that had been washed three times with distilled H_2_O and 100% isopropanol and dried under a stream of pressurized air. Gaskets were assembled on microscope coverslips and placed on the stage of a TwoMP mass photometer (MP, Refeyn Ltd, Oxford, UK) with immersion oil. Samples were measured in 1x phosphate-buffered saline (PBS, 10 mM Na_2_HPO_4_, 1.8 mM KH_2_PO_4_, 137 mM NaCl, 2.7 mM KCl (pH 7.4)). To this end, 18 µL 1x PBS was used to focus the MP before 2 µL sample (0.5 µM protein) was added, rapidly mixed, and measured.

Samples were prepared by diluting purified protein to 20 µM monomer concentration in desalting buffer (25 mM Tricine (pH 8.00), 75 mM NaCl), as determined by absorption at 280 nm. Immediately prior to measuring, samples were further diluted to a final concentration of 0.5 µM. For samples containing purified SSU, a 5-fold SSU excess was added to the undiluted sample shortly before dilution to 0.5 µM Rubisco monomer concentration.

Data was acquired for 60 s at 100 frames per second using AcquireMP (Refeyn Ltd, Oxford, UK). MP contrast was calibrated to molecular masses using 50 nM of an in-house purified protein mixture containing complexes of known molecular mass. MP datasets were processed and analyzed using DiscoverMP (Refeyn Ltd, Oxford, UK). Details of MP image analysis have been described previously^39^.

### Rubisco inhibition assays

To assess inhibition of Rubisco variants by RuBP, XuBP or CABP, 20 µM unactivated Rubiscos were incubated with 50 mM HEPES-KOH (pH 8.00), 4 mM EDTA, and either 3 mM RuBP, 1 mM XuBP, or 0.1 mM CABP for 30 minutes. Subsequently, inhibited Rubiscos were 1:10 diluted into standard photospectrometric assay reactions containing 100 mM HEPES-KOH (pH 8.00), 0.3 mM NADH, 10 mM MgCl_2_, 2.5 U/mL pgk, 5 U/mL gapdh, 5 mM ATP, 0.02 mg/mL carbonic anhydrase, 70 mM NaHCO_3_, and 3 mM RuBP. Reaction progress was followed via quantifying the decrease in absorption at 340 nm.

To assess the positive effects XuBP or CABP on activation, activation mixtures (50 mM HEPES-KOH (pH 8.00), 10 mM MgCl_2_, 20 mM NaHCO_3_, 0.02 mg/mL carbonic anhydrase, 20 µM Rubisco) were supplemented with 1 mM XuBP or 0.1 mM CABP, prior to 1:10 dilution into the aforementioned assay mixtures.

### Crystallization and structure determination

The sitting-drop vapor-diffusion method was used for crystallization at 16 °C. Purified AncL+7 r269W e271R l273N (10 mg/mL) was incubated with 300 µM CABP and 4.8 mM MgCl_2_ for 1 hour at 3% (v/v) CO_2_, prior to 1:1 mixing with 100 mM TRIS-HCl (pH 8.5), 200 mM MgCl_2_, 30 % (v/v) polyethylene glycol 400. Crystals appeared within 2 days and were flash frozen in liquid nitrogen.

For the co-crysallization of AncL+7 r269W e271R l273N with AncSSU, purified Rubisco was incubated with a 4-fold molar excess of purified AncSSU at 25 °C for 30 min, prior to purification of the L8S8 complex by size exclusion chromatography (see protein purification section).

Reconstituted L8S8 complex (9 mg/mL) was incubated with 350 µM CABP and 5.6 mM MgCl_2_ for 1 hour at 3% (v/v) CO_2_, prior to 1:1 mixing with 200 mM BIS-TRIS propane (pH 9.1), and 20 % (w/v) polyethylene glycol 4000. Crystals appeared within 2 days. The mother liquor was supplemented with 25 % (v/v) PEG200 before crystals were flash frozen in liquid nitrogen.

X-ray diffraction data (Extened Data Table 1) were collected at the beamline PETRA III P14 of the DESY (Deutsches Elektronen-Synchrotron, Hamburg). Data were processed with the XDS software package^40^. Structures were solved by molecular replacement using Phaser of the Phenix software package^41^ (v.1.1.14), and refined with Phenix.Refine. Additional modelling, manual refinement, and ligand fitting was done in Coot^42^ (v.0.9.8.3). Final positional and B-factor refinements, as well as water picking, were performed using Phenix.Refine. Structural models for the L8 and L8S8 complex of AncL+7 r269W e271R l273N were deposited to the Protein Data Bank in Europe (PDBe) under PDB accession 8QMV and 8QMW, respectively. Figures were made using PyMOL 2.4.1.

### Hydrogen-deuterium exchange (HDX) mass spectrometry

HDX-MS experiments on Rubisco were carried out as described previously^43^ with minor modifications. The conformational dynamics of Rubisco’s LSU was studied in 2 different states, i.e., LSU and LSU/CABP (see **Source Data HDX**). Depending on the investigated state, LSU and SSU were either used individual or mixed to reach final protein concentrations of 25 µM. Where indicated, CABP was present at 50 µM. These batch solutions were incubated in 3% (v/v) CO_2_ atmosphere for 60 min at 30 °C to facilitate carbamylation of AncL+7 residue 187^9^. The protein batch solutions were stored in a cooled tray (1 °C) from which 7.5 μl were withdrawn by a robotic autosampler unit (LEAP technologies) and mixed with 67.5 μl of D_2_O-containing buffer (20 mM Tricine-Na pH 8.0, 75 mM NaCl) to start an HDX reaction. After 10, 30, 95, 1,000 or 10,000 s of incubation in another tray at 25 °C, 55 μl samples were taken from the reaction and mixed with an equal volume of pre-dispensed quench buffer (400 mM KH_2_PO_4_/H_3_PO_4_, 2 M guanidine-HCl, pH 2.2) kept at 1 °C. 95 µl of the resulting mixture were injected through a 50 µl sample loop into an ACQUITY UPLC M-Class System with HDX Technology (Waters)^44^. Undeuterated samples were generated and treated similar except that H_2_O-containing buffer was employed for dilution followed by incubation for 10 s. The injected samples were flushed out of the loop with H_2_O + 0.1% (v/v) formic acid at 100 µl/min flow rate, guided to an Enzymate BEH Pepsin Column (5 µm, 2.1 x 30 mm (Waters)) containing immobilized porcine pepsin for proteolytic digestion of the proteins at 12 °C, and the resulting peptic peptides collected on an ACQUITY UPLC BEH C18 VanGuard Pre-column (1.7 µm, 2.1 mm x 5 mm (Waters)) kept at 0.5 °C. After 3 min of digestion and trapping, the trap column was placed in line with an ACQUITY UPLC BEH C18 column (1.7 μm, 1.0 x 100 mm (Waters)), and the peptides were eluted at 0.5 °C using a gradient of H_2_O + 0.1% (v/v) formic acid (eluent A) and acetonitrile + 0.1% (v/v) formic acid (eluent B) at a flow rate of 30 μl/min as follows: 0-7 min/95-65% A, 7-8 min/65-15% A, 8-10 min/15% A, 10-11 min/5% A, 11-16 min/95% A. The peptides were ionized with an electrospray ionization source (250 °C capillary temperature, 3.0 kV spray voltage), and mass spectra were acquired in positive ion mode over a range of 50 to 2000 *m/z* on a G2-Si HDMS mass spectrometer with ion mobility separation (Waters) using Enhanced High Definition MS (HDMS^E^) or High Definition MS (HDMS) mode for undeuterated and deuterated samples, respectively^45^. Lock mass correction was implemented with [Glu1]-Fibrinopeptide B standard (Waters). During separation of the peptides, the protease column was washed three times by injecting 80 µl of 0.5 M guanidine-HCl in 4% (v/v) acetonitrile. Additionally, blank injections were performed between each sample to minimize peptide carry-over. All measurements were carried out in three technical replicates.

Peptides were identified from the undeuterated samples (acquired with HDMS^E^) with search parameters as described previously^43^ with the software ProteinLynx Global SERVER (PLGS, Waters), using low energy, elevated energy, and intensity thresholds of 300, 100 and 1,000 counts, respectively, and matched using a database containing the amino acid sequences of LSU, SSU, porcine pepsin, and their reversed sequence. After automated processing of the spectra with the software DynamX (Waters), all spectra were inspected manually and, if necessary, peptides were omitted (e.g. in case of a low signal-to-noise ratio or the presence of overlapping peptides).

## Extended Data

**Extended Data Figure 1.**
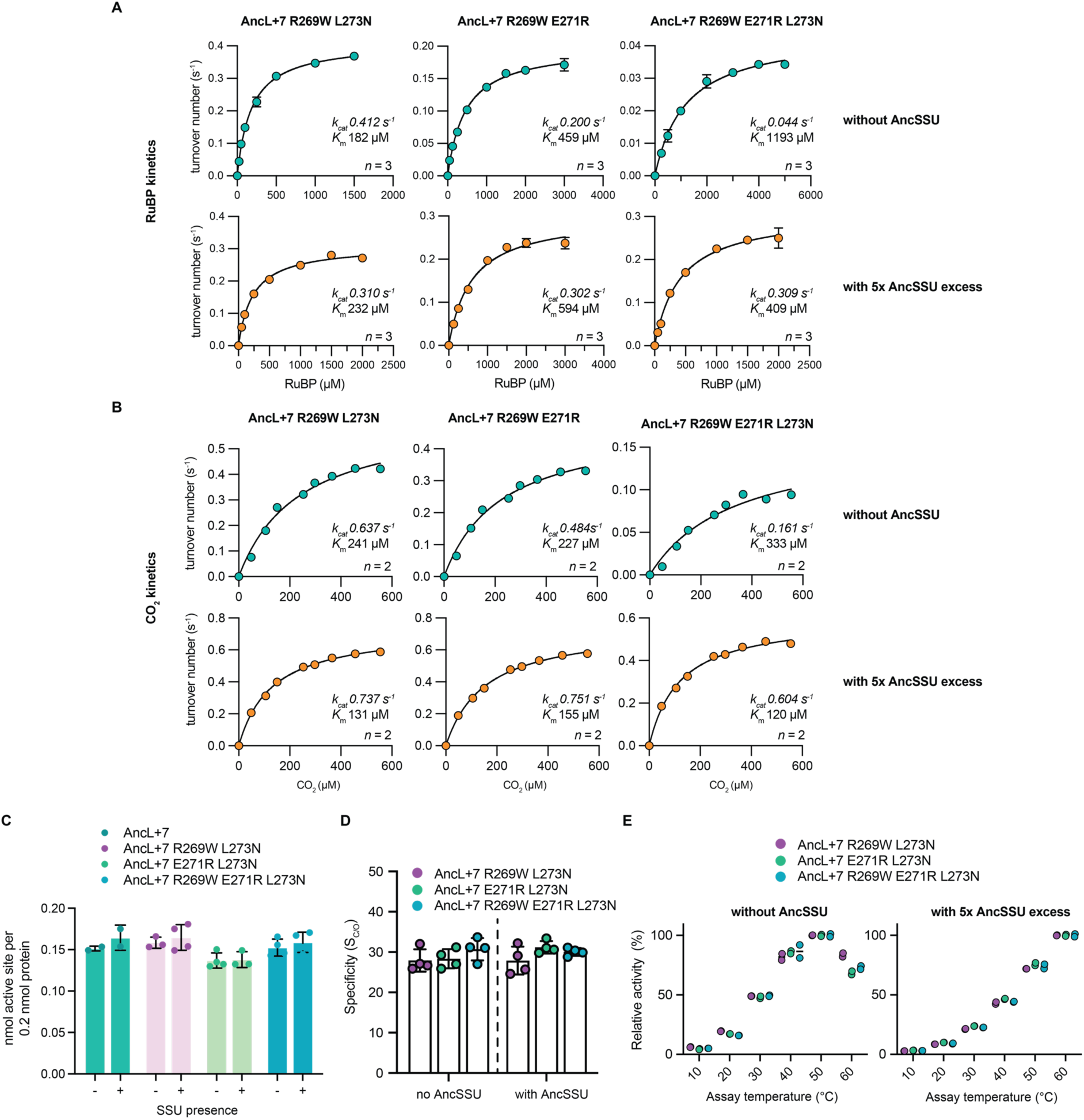
(**A**) RuBP kinetics of AncL+7-based variants AncL+7 R269W L273N, AncL+7 R269W E271R, and R269W E271R L273N measured without (teal) or with (orange) a 5x AncSSU excess. *N* = 3. *k*cat and *K*m are listed in the plots. (**B**) CO2 kinetics of AncL+7-based variants AncL+7 R269W L273N, AncL+7 R269W E271R, and R269W E271R L273N measured without (teal) or with (orange) a 5x AncSSU excess. *N* = 3. *k*cat and *K*m are listed in the plots. (**C**) Relative CABP binding of relevant Rubisco variants with or without presence of a 5-fold AncSSU excess. (**D**) Specificity (SC/O) of relevant Rubisco variants with or without presence of a 5-fold AncSSU excess. (**E**) Relative activity of relevant Rubisco variants at varying temperatures without AncSSU (left) and with a 5-fold AncSSU excess (right). Activity is relative to the highest measured activity of the variant in question, for either the dataset with or without AncSSU.

**Extended Data Figure 2.**
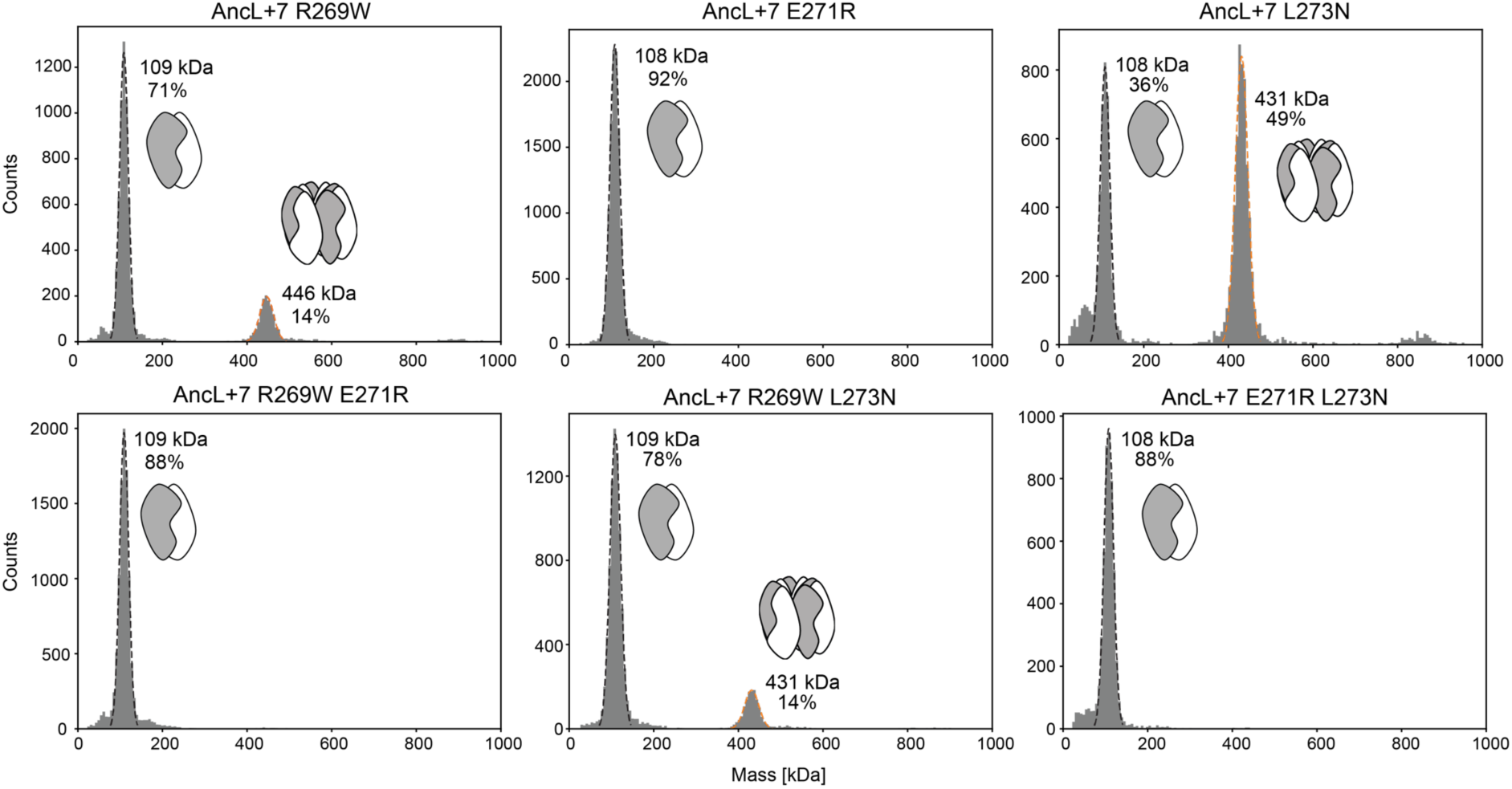
Mass photometry characterization of AncL+7-based single and double substitution constructs on the trajectory to the AncL+7 R269W E271R L273N. Single substitution constructs in top row, double substitution constructs in bottom row. Percentages indicate integrated counts relative to all counts of the measurement.

**Extended Data Figure 3.**
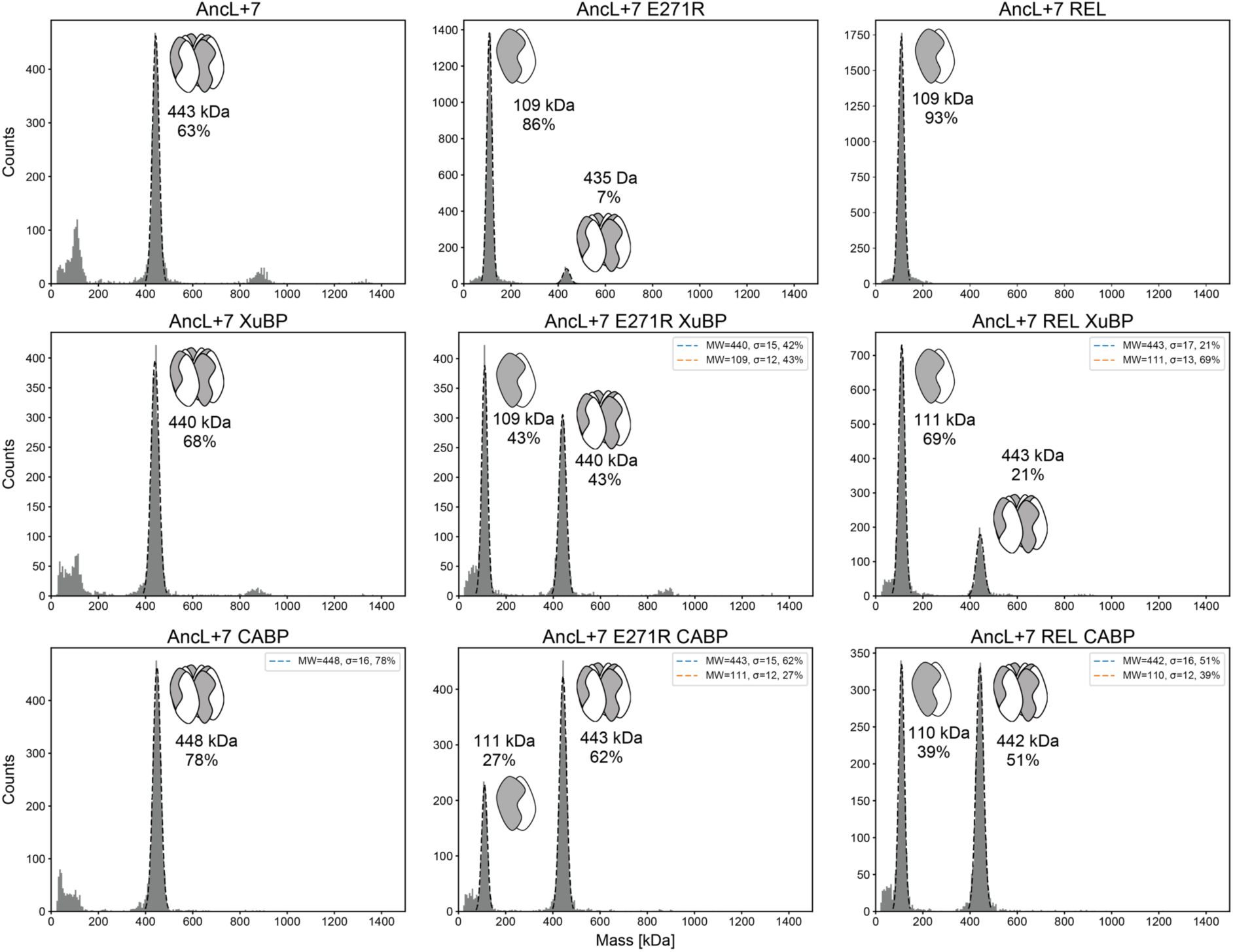
Mass photometry characterization of AncL+7, AncL+7 E271R, and AncL+7 R269W E271R L273N (denoted as AncL+7 REL) alone, with xylulose-1,5-bisphosphate (XuBP, 1 mM) and carboxyarabinitol-bisphosphate (CABP, 0.1 mM). First row contains measurements in isolation, second row contains measurements with XuBP, third row contains measurements with CABP. Percentages indicate integrated counts relative to all counts of the measurement.

**Extended Data Figure 4.**
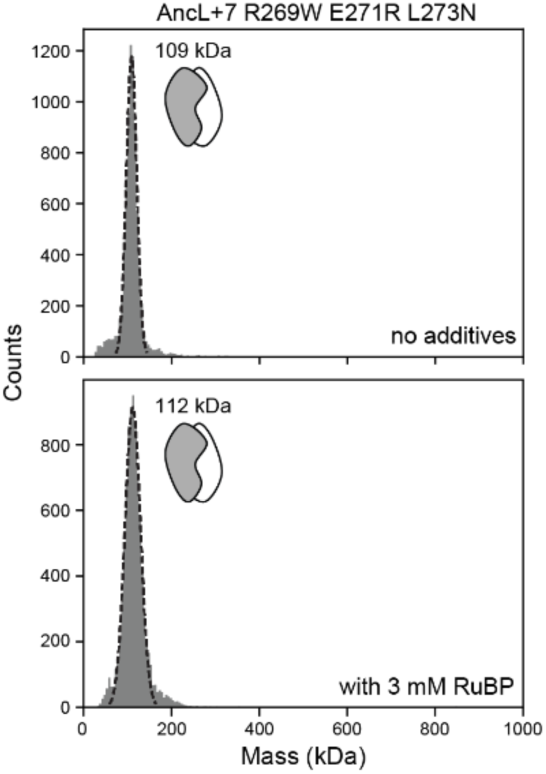
Mass photometry characterization of AncL+7 R269W E271R L273N without additives and in presence of 3 mM RuBP. Top shows measurement in isolation, bottom shows measurements with RuBP. Protein was pre-incubated at 20 µM concentration in presence of 3 mM RuBP for >30 min prior to crash dilution to 500 nM protein concentration and subsequent measurement at 50 nM protein concentration. Theoretical mass of a dimer is 107 kDa and of an octamer 428 kDa.

**Extended Data Figure 5.**
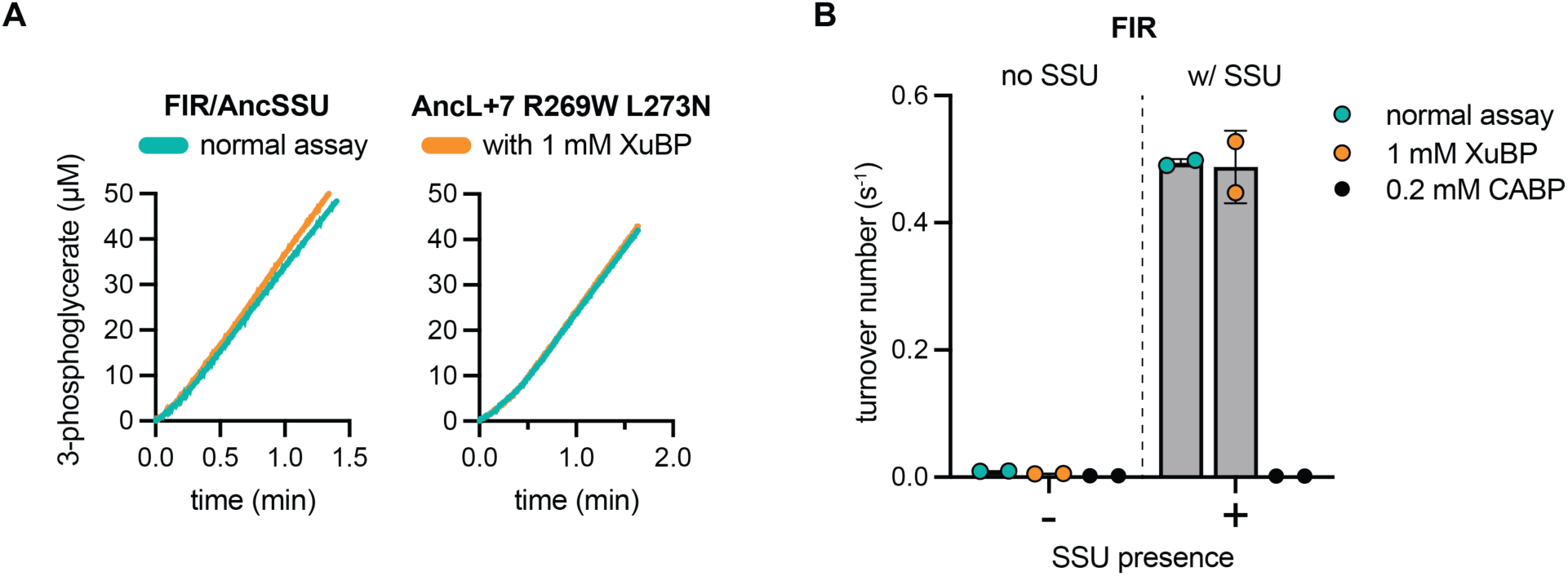
AncL-based variants are not inhibited by XuBP. (**A**) Unchanged 3-phosphoglycerate production of FIR construct or AncL+7 R269W L273N either in isolation or in presence of 1 mM XuBP. (**B**) Absolute turnover number (s^-1^) of FIR construct shows SSU dependence and unchanged turnover in presence of 1 mM XuBP, while the addition of 0.2 mM CABP fully inhibits the enzyme.

**Extended Data Figure 6.**
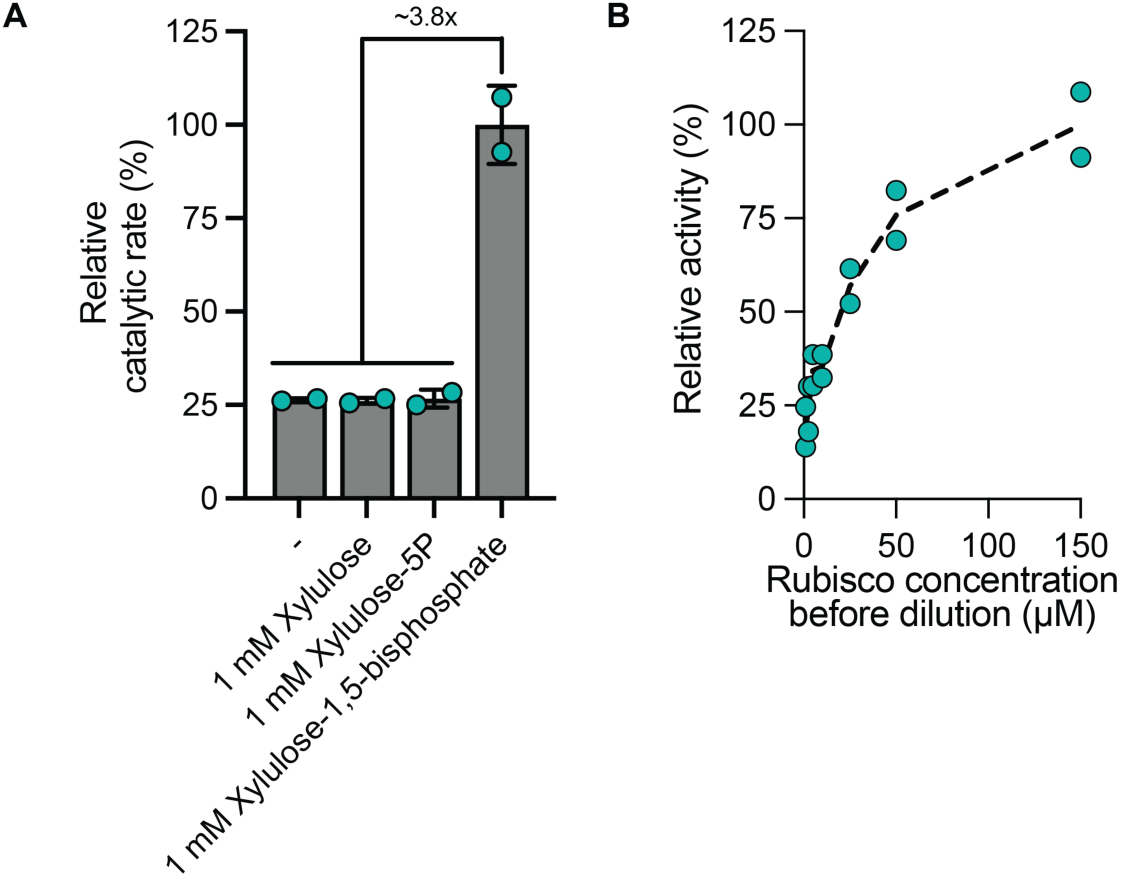
**(A)** Activity of AncL+7 R269W, E271R, L273N only increases in presence of 1 mM xylulose-1,5-bisphosphate and is unresponsive to the addition of mono- or non-phosphorylated xylulose. **(B)** Activity of AncL+7 REL crash diluted into assay mixture (normalized to the same concentration in the assay) from different concentrations shows a positive correlation with increasing pre-dilution concentrations. Activity is given relative to the measurements of 150 µM Rubisco pre-dilution.

**Extended Data Figure 7.**
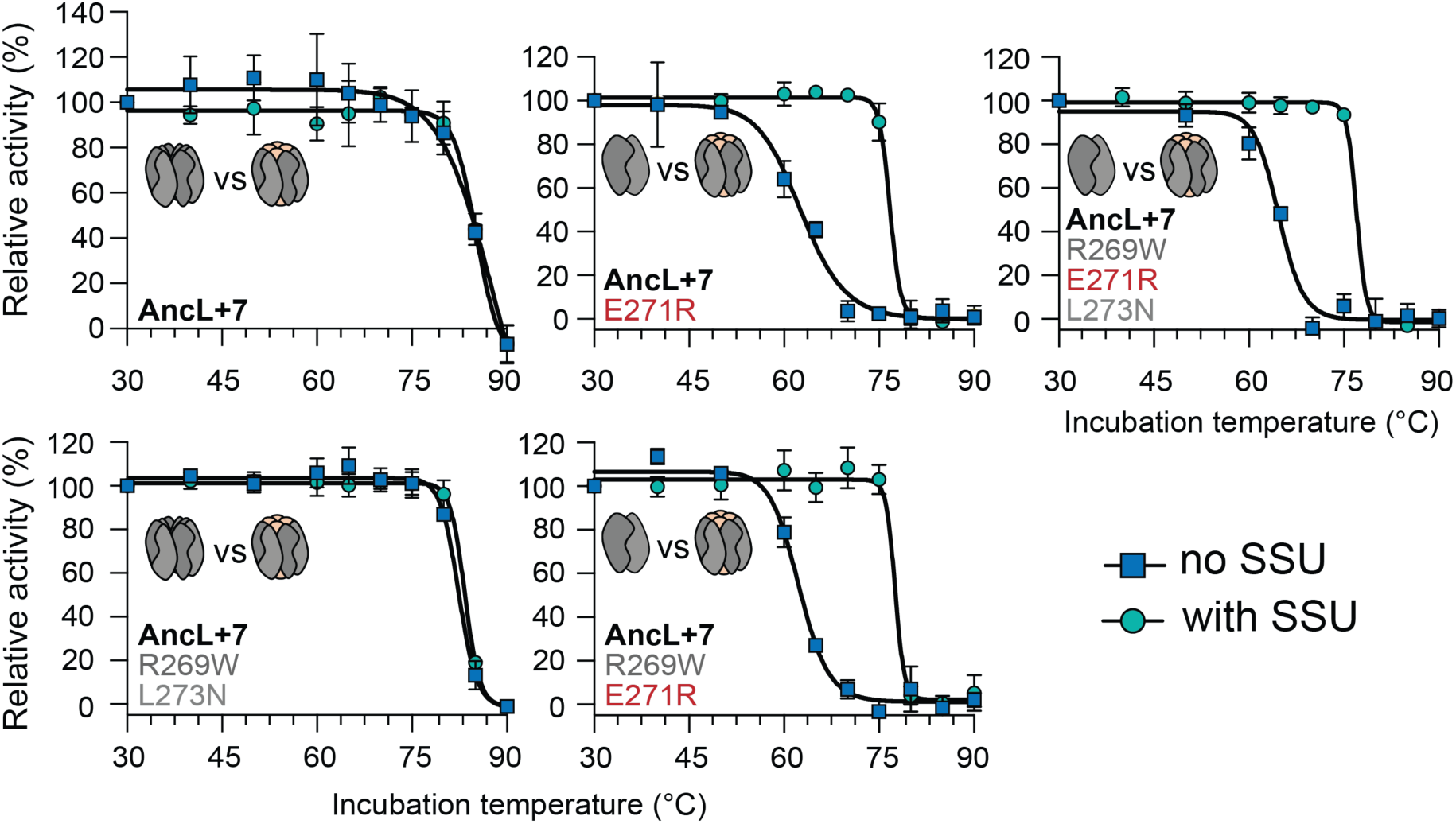
Denaturation of AncL+7-based variants on the trajectory to AncL+7 R269W E271R L273N indicate that variants which contain E271R (highlighted in red) and thus disassemble into dimers in absence of AncSSU are destabilized to thermal denaturation (see cartoon schemes for inferred oligomeric state). Variants were incubated at indicated temperatures for 1 hour prior to cooling down and measuring remaining activity at 25 °C. A 5-fold AncSSU excess was used for incubations in presence of SSU.

**Extended Data Figure 8.**
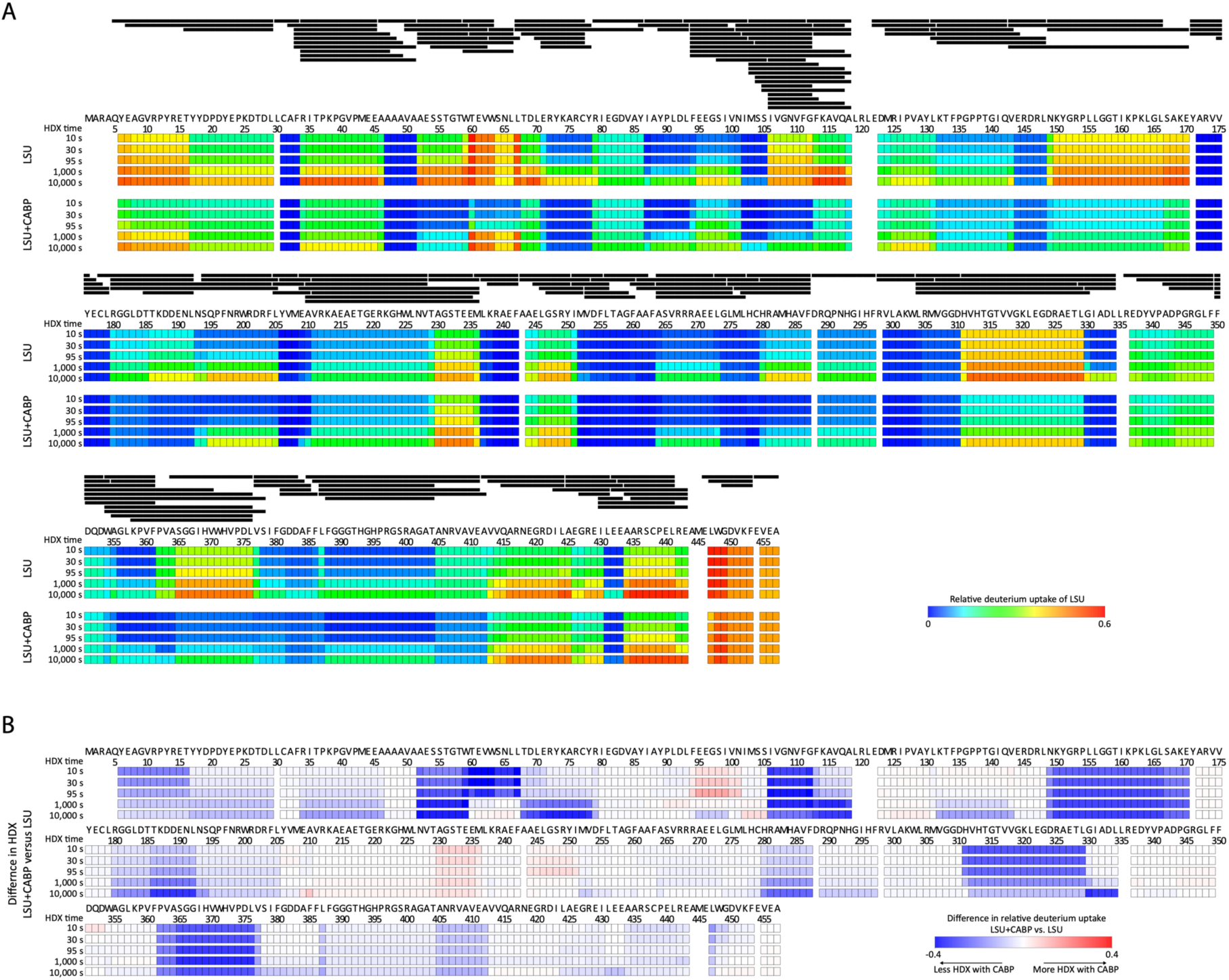
Hydrogen-deuterium exchange mass spectrometry of AncL+7 LSU and LSU+CABP. (**A**) Relative deuterium uptake of Rubisco’s large subunit in LSU and LSU+CABP conditions. Detected peptides are indicated as black bars above the measurements. (**B**) Difference in hydrogen-deuterium exchange between LSU+CABP versus just the LSU.

**Extended Data Table 1.**

| Rubisco variant | AncL+7 REL<br>(PDB 8QMV) | AncL+7 REL<br>AncSSU<br>(PDB 8QMW) |
| --- | --- | --- |
| **Data collection** |  |  |
| Beamline | PETRA III P14 | PETRA III P14 |
| Wavelength (Å) | 0.6888 | 0.6888 |
| Space Group | C 2 2 2<sub>1</sub> | C 1 2 1 |
| Unit cell dimensions |  |  |
| a, b, c (Å) | 121.79, 203.80, 147.37 | 206.01, 106.44, 108.49 |
| α, β, γ (°) | 90.00, 90.00, 90.00 | 90.00, 113.09, 90.00 |
| Resolution (Å) | 24.97 - 1.85 (1.916 - 1.850) | 24.88 - 1.75 (1.813 - 1.750) |
| Unique reflections | 155284 (15351) | 213396 (21212) |
| Multiplicity | 13.9 (12.6) | 7.2 (7.3) |
| Completeness (%) | 99.9 (99.6) | 98.6 (98.4) |
| *I* / $\sigma$ | 16.0 (2.2) | 11.1 (2.7) |
| *R*<sub>merge</sub> | 0.113 (1.04) | 0.124 (0.653) |
| *R*<sub>pim</sub> | 0.031 (0.302) | 0.049 (0.258) |
| CC<sub>1/2</sub> | 0.999 (0.762) | 0.997 (0.834) |
| **Refinement** |  |  |
| *R*<sub>work</sub> / *R*<sub>free</sub> | 0.1607 / 0.1739 | 0.1448 / 0.1745 |
| RMS bonds | 0.003 | 0.013 |
| RMS angles | 0.666 | 1.148 |
| Ramachandran |  |  |
| favored (%) | 97.30 | 96.56 |
| allowed (%) | 2.70 | 3.44 |
| outliers (%) | 0.00 | 0.00 |
| Number of atoms | 15421 | 19717 |
| Protein | 14180 | 17595 |
| Ligands | 88 | 107 |
| Solvent | 1153 | 2015 |
| Average B-factor | 28.46 | 18.47 |
| Protein | 28.05 | 17.46 |
| Ligands | 23.02 | 15.25 |
| Solvent | 33.84 | 27.42 |

Values in parentheses are for highest-resolution shell.

**Supplementary Table 1.**
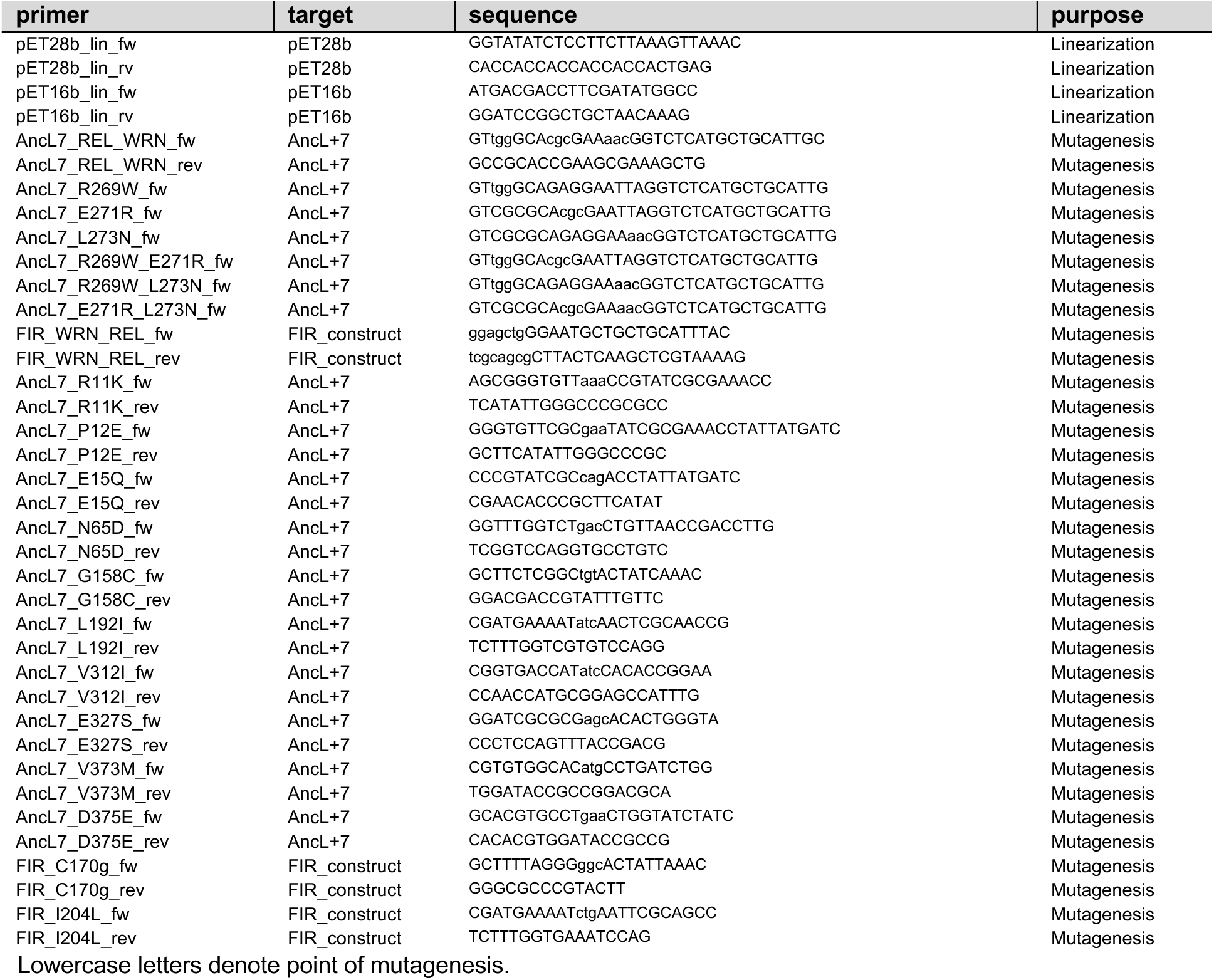
List of primers used in this study.

**Supplementary Table 2.** List of plasmids used in this study.

| plasmid | relevant features | source or reference |
| --- | --- | --- |
| pET16b | *E. coli* expression vector, T7 promoter, Amp<sup>R</sup>, *lacI*, N-terminal His<sub>10</sub>-tag | Merck Chemicals |
| pET28b | *E. coli* expression vector, T7 promoter, Kan<sup>R</sup>, *lacI*, C-terminal His<sub>6</sub>-tag | Merck Chemicals |
| pET16b-AncL+7 | *E. coli* expression vector, T7 promoter, Amp<sup>R</sup>, *lacI*, N-terminal His<sub>10</sub>-tag | (ref. <sup>9</sup>) |
| pET16b-fiber interface reversion | *E. coli* expression vector, T7 promoter, Amp<sup>R</sup>, *lacI*, N-terminal His<sub>10</sub>-tag | (ref. <sup>9</sup>) |

Genes cloned into pET16b vectors were cloned in-frame with an N-terminal His-tag encoding sequence on the LSU. Given list does not include point mutant variant-encoding plasmids derived from the plasmid above via Q5 mutagenesis.

**Supplementary Table 3.**
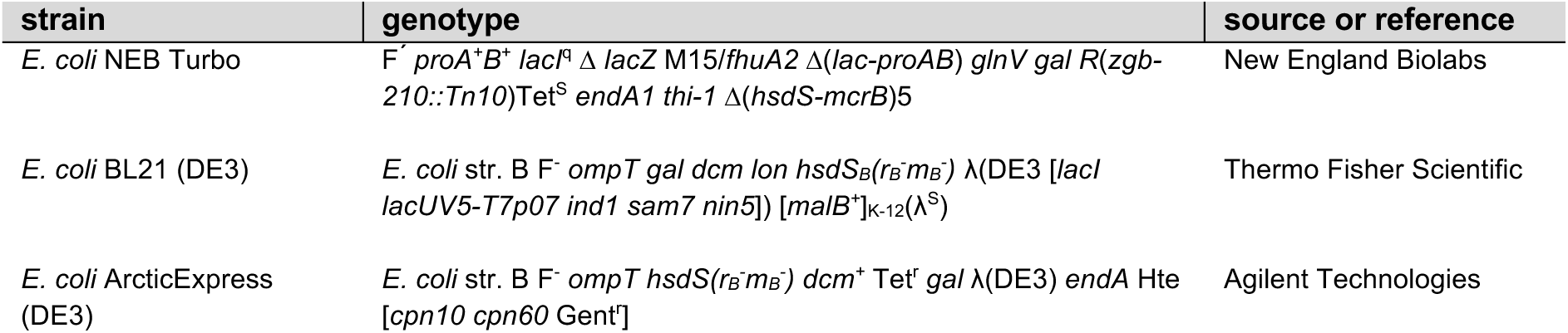
List of strains used in this study.

## References

1. Muller, H. J. Genetic variability, twin hybrids and constant hybrids, in a case of balanced lethal factors. Genetics 3, (1918).

2. Ali, M. H. & Imperiali, B. Protein oligomerization: How and why. Bioorg Med Chem 13, 5013–5020 (2005).

3. Hochberg, G. K. A. et al. A hydrophobic ratchet entrenches molecular complexes. Nature 588, (2020).

4. Schulz, L., Sendker, F. L. & Hochberg, G. K. A. Non-adaptive complexity and biochemical function. Curr Opin Struct Biol 73, 102339 (2022).

5. Force, A. et al. Preservation of duplicate genes by complementary, degenerative mutations. Genetics 151, 1531–45 (1999).

6. Stoltzfus, A. On the possibility of constructive neutral evolution. J Mol Evol 49, 169– 181 (1999).

7. Stoltzfus, A. On the possibility of constructive neutral evolution. J Mol Evol (1999) doi:10.1007/PL00006540.

8. Muñoz-Gómez, S. A., Bilolikar, G., Wideman, J. G. & Geiler-Samerotte, K. Constructive neutral evolution 20 years later. J Mol Evol 89, 172–182 (2021).

9. Schulz, L. et al. Evolution of increased complexity and specificity at the dawn of form I Rubiscos. Science *(1979)* 378, 155–160 (2022).

10. Emlaw, J. R. et al. A single historical substitution drives an increase in acetylcholine receptor complexity. Proc Natl Acad Sci U S A (2021) doi:10.1073/pnas.2018731118.

11. Finnigan, G. C., Hanson-Smith, V., Stevens, T. H. & Thornton, J. W. Evolution of increased complexity in a molecular machine. Nature 481, (2012).

12. Andrews, T. J. Catalysis by cyanobacterial ribulose-bisphosphate carboxylase large subunits in the complete absence of small subunits. J Biol Chem (1988).

13. Liu, C. et al. Coupled chaperone action in folding and assembly of hexadecameric Rubisco. Nature (2010) doi:10.1038/nature08651.

14. Joshi, J., Mueller-Cajar, O., Tsai, Y.-C. C., Hartl, F. U. & Hayer-Hartl, M. Role of small subunit in mediating assembly of red-type Form I Rubisco. Journal of Biological Chemistry 290, 1066–1074 (2015).

15. Spreitzer, R. J. Role of the small subunit in ribulose-1,5-bisphosphate carboxylase/oxygenase. Arch Biochem Biophys 414, 141–149 (2003).

16. Garcia-Seisdedos, H., Empereur-Mot, C., Elad, N. & Levy, E. D. Proteins evolve on the edge of supramolecular self-assembly. Nature (2017) doi:10.1038/nature23320.

17. Liu, A. K. et al. Structural plasticity enables evolution and innovation of RuBisCO assemblies. Sci Adv 8, (2022).

18. Gunn, L. H., Valegard, K. & Andersson, I. A unique structural domain in Methanococcoides burtonii ribulose-1,5-bisphosphate carboxylase/oxygenase (Rubisco) acts as a small subunit mimic. Journal of Biological Chemistry 292, (2017).

19. Bracher, A., Sharma, A., Starling-Windhof, A., Hartl, F. U. & Hayer-Hartl, M. Degradation of potent Rubisco inhibitor by selective sugar phosphatase. Nat Plants 1, (2015).

20. Parry, M. A. J., Keys, A. J., Madgwick, P. J., Carmo-Silva, A. E. & Andralojc, P. J. Rubisco regulation: a role for inhibitors. J Exp Bot 59, 1569–1580 (2007).

21. Taverna, D. M. & Goldstein, R. A. Why are proteins marginally stable? *Proteins: Structure*, Function and Genetics 46, (2002).

22. Tokuriki, N. & Tawfik, D. S. Chaperonin overexpression promotes genetic variation and enzyme evolution. Nature (2009) doi:10.1038/nature08009.

23. Bhattacharya, S. et al. NMR-guided directed evolution. Nature 610, (2022).

24. Andersson, I. & Backlund, A. Structure and function of Rubisco. Plant Physiology and Biochemistry (2008) doi:10.1016/j.plaphy.2008.01.001.

25. Taylor, T. C. & Andersson, I. Structural transitions during activation and ligand binding in hexadecameric Rubisco inferred from the crystal structure of the activated unliganded spinach enzyme. Nat Struct Biol 3, (1996).

26. Andrews, T. J. Catalysis by cyanobacterial ribulose-bisphosphate carboxylase large subunits in the complete absence of small subunits. Journal of Biological Chemistry 263, 12213–12219 (1988).

27. Aigner, H. et al. Plant RuBisCo assembly in E. coli with five chloroplast chaperones including BSD2. Science *(1979)* 358, (2017).

28. Portis, A. R. Rubisco activase - Rubisco’s catalytic chaperone. Photosynthesis Research vol. 75 Preprint at DOI:10.1023/A:1022458108678 (2003).

29. Portis, A. R. The regulation of Rubisco by Rubisco activase. J Exp Bot 46, (1995).

30. Tokuriki, N. & Tawfik, D. S. Chaperonin overexpression promotes genetic variation and enzyme evolution. Nature 459, 668–673 (2009).

31. Pierce, J., Tolbert, N. E. & Barker, R. Interaction of Ribulosebisphosphate Carboxylase/Oxygenase with Transition-State Analogs. Biochemistry (1980) doi:10.1021/bi00546a018.

32. Kane, H. et al. An improved method for measuring the CO2/O2 specificity of ribulosebisphosphate carboxylase-oxygenase. Functional Plant Biology 21, 449 (1994).

33. Bracher, A., Sharma, A., Starling-Windhof, A., Hartl, F. U. & Hayer-Hartl, M. Degradation of potent Rubisco inhibitor by selective sugar phosphatase. Nat Plants 1, (2015).

34. Gibson, D. G. et al. Enzymatic assembly of DNA molecules up to several hundred kilobases. Nat Methods (2009) doi:10.1038/nmeth.1318.

35. Karkehabadi, S. et al. Chimeric small subunits influence catalysis without causing global conformational changes in the crystal structure of ribulose-1,5-bisphosphate carboxylase/oxygenase. Biochemistry 44, 9851–9861 (2005).

36. Joshi, J., Mueller-Cajar, O., Tsai, Y.-C. C., Hartl, F. U. & Hayer-Hartl, M. Role of small subunit in mediating assembly of red-type Form I Rubisco. Journal of Biological Chemistry 290, 1066–1074 (2015).

37. Wilson, R. H., Martin-Avila, E., Conlan, C. & Whitney, S. M. An improved *Escherichia coli* screen for Rubisco identifies a protein–protein interface that can enhance CO2-fixation kinetics. Journal of Biological Chemistry 293, 18–27 (2018).

38. Kubien, D. S., Brown, C. M. & Kane, H. J. Quantifying the amount and activity of Rubisco in leaves. in *Methods in molecular biology (Clifton*, N.J.) 349–362 (2011). doi:10.1007/978-1-60761-925-3_27.

39. Sonn-Segev, A. et al. Quantifying the heterogeneity of macromolecular machines by mass photometry. Nat Commun (2020) doi:10.1038/s41467-020-15642-w.

40. Kabsch, W. XDS. Acta Crystallogr D Biol Crystallogr 66, 125–132 (2010).

41. Adams, P. D. et al. PHENIX: a comprehensive Python-based system for macromolecular structure solution. Acta Crystallogr D Biol Crystallogr 66, 213–221 (2010).

42. Emsley, P. & Cowtan, K. Coot: model-building tools for molecular graphics. Acta Crystallogr D Biol Crystallogr 60, 2126–2132 (2004).

43. Osorio-Valeriano, M. et al. ParB-type DNA segregation proteins are CTP-dependent molecular switches. Cell 179, 1512–1524.e15 (2019).

44. Wales, T. E., Fadgen, K. E., Gerhardt, G. C. & Engen, J. R. High-speed and high-resolution UPLC separation at zero degrees celsius. Anal Chem 80, 6815–6820 (2008).

45. Geromanos, S. J. et al. The detection, correlation, and comparison of peptide precursor and product ions from data independent LC-MS with data dependant LC-MS/MS. Proteomics 9, 1683–1695 (2009).

